# Deep Learning-Based Point-Scanning Super-Resolution Imaging

**DOI:** 10.1101/740548

**Authors:** Linjing Fang, Fred Monroe, Sammy Weiser Novak, Lyndsey Kirk, Cara R. Schiavon, Seungyoon B. Yu, Tong Zhang, Melissa Wu, Kyle Kastner, Yoshiyuki Kubota, Zhao Zhang, Gulcin Pekkurnaz, John Mendenhall, Kristen Harris, Jeremy Howard, Uri Manor

## Abstract

Point scanning imaging systems (e.g. scanning electron or laser scanning confocal microscopes) are perhaps the most widely used tools for high resolution cellular and tissue imaging. Like all other imaging modalities, the resolution, speed, sample preservation, and signal-to-noise ratio (SNR) of point scanning systems are difficult to optimize simultaneously. In particular, point scanning systems are uniquely constrained by an inverse relationship between imaging speed and pixel resolution. Here we show these limitations can be mitigated via the use of deep learning-based super-sampling of undersampled images acquired on a point-scanning system, which we termed point-scanning super-resolution (PSSR) imaging. Oversampled, high SNR ground truth images acquired on scanning electron or Airyscan laser scanning confocal microscopes were ‘crappified’ to generate semi-synthetic training data for PSSR models that were then used to restore real-world undersampled images. Remarkably, our EM PSSR model could restore undersampled images acquired with different optics, detectors, samples, or sample preparation methods in other labs. PSSR enabled previously unattainable 2 nm resolution images with our serial block face scanning electron microscope system. For fluorescence, we show that undersampled confocal images combined with a multiframe PSSR model trained on Airyscan timelapses facilitates Airyscan-equivalent spatial resolution and SNR with ∼100x lower laser dose and 16x higher frame rates than corresponding high-resolution acquisitions. In conclusion, PSSR facilitates point-scanning image acquisition with otherwise unattainable resolution, speed, and sensitivity.

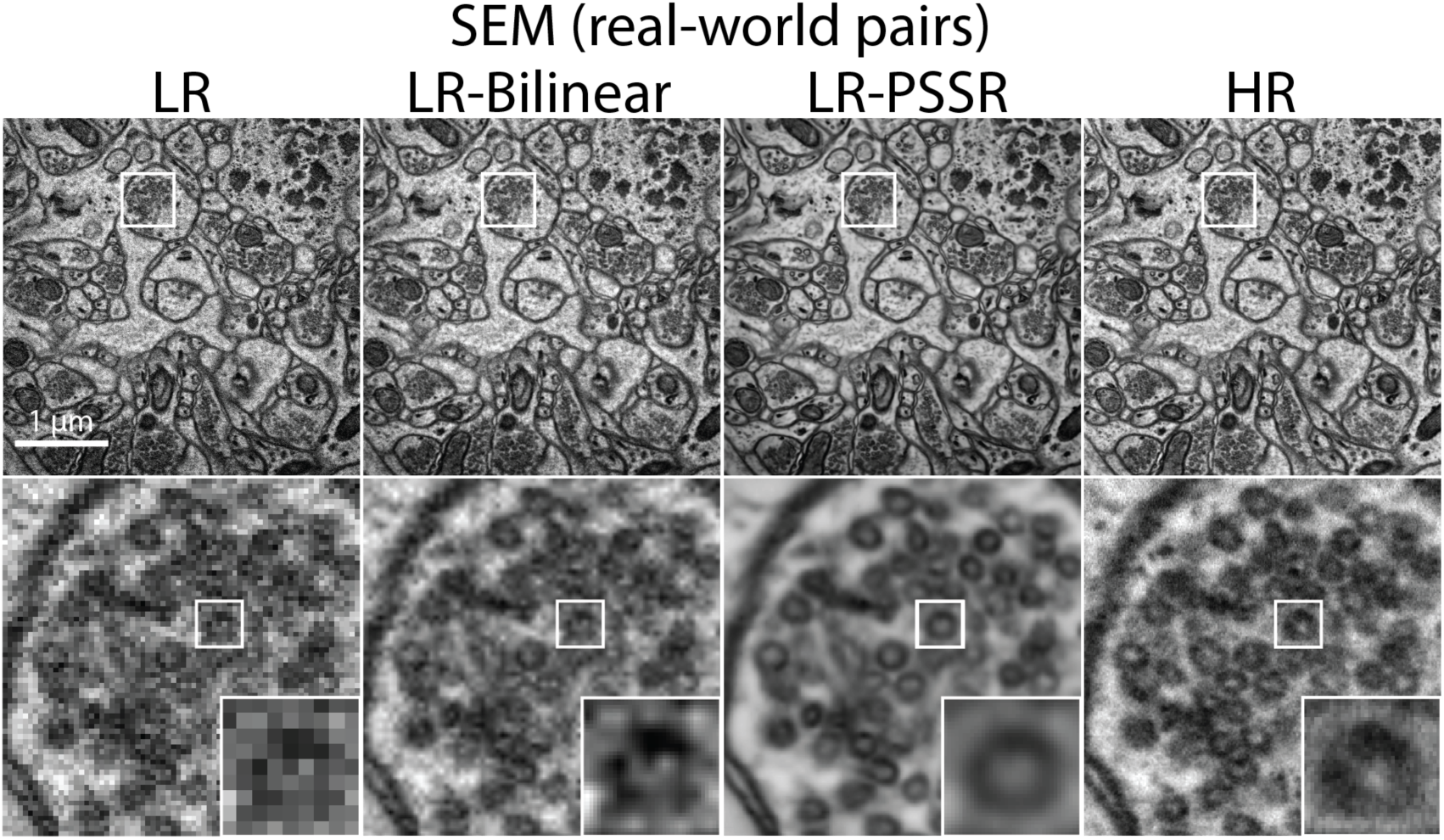

## Introduction

An essential tool for understanding the spatiotemporal organization of biological systems, microscopy is nearly synonymous with biology itself. Like all imaging systems, microscopes suffer from the so-called “eternal triangle of compromise”, which dictates that image resolution, illumination intensity (and therefore sample damage), and imaging speed are all in tension with one another. In other words, it is usually impossible to optimize one parameter without compromising at least one of the others. This is particularly noticeable for point-scanning systems, e.g. scanning electron (SEM) and laser scanning confocal (LSM) microscopes, for which higher resolution images require higher numbers of sequentially acquired pixels to ensure proper sampling, thus increasing the imaging time and sample damage in direct proportion to the pixel resolution. In spite of these limitations, point-scanning systems remain perhaps the most common imaging modality in biological research labs due to their versatility and ease of use for a broad range of applications.

“Super-resolution” deep learning has been extensively used to “super-sample” the pixels in down-sampled digital images, effectively increasing their resolution^1-3^. For microscopy, deep learning has long been established as an optimal method for image analysis and segmentation^4^. More recently, deep learning has been employed with spectacular results in restoring microscopy images from relatively noisy, low resolution acquisitions to high resolution outputs that have a high signal-to-noise ratio (SNR)^5-13^. Similarly, anisotropic fluorescence and EM volumetric data has been supersampled to isotropy using deep learning^5,14^. Here we show that deep learning-based restoration of 16x undersampled images facilitates faster, lower dose imaging on both SEM and scanning confocal microscopes, which in turn allows for 16x higher imaging speeds, ≥16x lower sample damage, and 16x smaller raw image file sizes due to the 16x smaller number of pixels acquired. Thus, the <u>P</u>oint <u>S</u>canning <u>S</u>uper-<u>R</u>esolution (PSSR) approach provides a strategy for increasing the spatiotemporal resolution of point scanning imaging systems to previously unattainable levels due to limitations imposed by sample damage or imaging speed when imaging at full pixel resolution.

## Results

Three-dimensional electron microscopy (3DEM) is a powerful technique for determining the volumetric ultrastructure of tissues, which is invaluable for connectomics research. In addition to serial section EM (ssEM)^15^ and focused ion beam SEM (FIB-SEM)^16^, one of the most common tools for high throughput 3DEM imaging is serial blockface scanning electron microscopy (SBFSEM)^17^, wherein a built-in ultramicrotome iteratively cuts ultrathin sections (usually between 50 - 100 nm) off the surface of a blockface after it was imaged with a scanning electron probe. This method facilitates relatively automated, high-throughput 3DEM imaging with minimal post-acquisition image alignment. Unfortunately, higher electron doses cause sample charging, which renders the sample too soft to section and image reliably (Supplementary Movie 1). Furthermore, the extremely long imaging times and large file sizes inherent to high resolution 3DEM imaging of relatively large volumes present a significant bottleneck for many labs. Thus, most 3DEM datasets are acquired with sub-Nyquist sampling (e.g. pixel sizes ≥ 4 nm), which precludes the reliable detection or analysis of smaller subcellular structures, such as ∼35 nm presynaptic vesicles. While undersampled 3DEM datasets can be suitable for many analyses, it would be useful to be able to mine targeted regions of these large datasets for higher resolution ultrastructural information. Unfortunately, many 3DEM imaging approaches are destructive, and high resolution ssEM can be slow and laborious. Thus, the ability to computationally increase the resolution of these datasets is of high value and broad utility.

Frustrated by our inability to perform SBFSEM imaging with the desired 2 nm resolution and SNR necessary to reliably detect presynaptic vesicles, we decided to test whether a deep convolutional neural net model (PSSR) trained on 2 nm high resolution (HR) images could “super-resolve” 8 nm low resolution (LR) images to 2 nm resolution (Fig. 1). To train a model for this purpose, many perfectly aligned high- and low-resolution image pairs are required. Instead of manually acquiring high- and low-resolution image pairs for training, we opted to generate semi-synthetic training data by computationally “crappifying” high-resolution images to simulate what their low-resolution counterparts might look like when acquired at the microscope. For this purpose we used ∼130 GB training data of 2 nm pixel transmission-mode scanning electron microscope (tSEM^18^) images of 40 nm ultrathin sections from the hippocampus of a Long Evans male rat. To generate semi-synthetic training pairs, we applied aggressive downsampling and degradation filters to our HR data, including heavy gaussian blur, random pixel shifts, and random salt-and-pepper noise in addition to 16x downsampling of the pixel resolution. We then trained our image pairs on a ResNet-based U-Net model (Fig. 1a – see Methods and Supplemental Tables for full details). Using a Mean Squared Error (MSE) loss function yielded excellent results as determined by visual inspection as well as Peak-Signal-to-Noise Ratio (PSNR), Structural Similarity (SSIM) measurements, and Fourier Ring Correlation (FRC) analysis. The PSSR-restored images from our semi-synthetic pairs contained more detail and yet displayed less noise, making it easier to discern fine details such as presynaptic vesicles (Fig. 1b). We next tested whether our PSSR model was effective on “real-world” LR images. Usually deep learning-based image restoration models are extremely sensitive to variations in image properties, precluding the use of a model generated from training images acquired in one condition on images acquired in another (i.e. data generated using a different sample preparation technique, type, or on a different microscope). As mentioned above, our training images were generated from 40 nm sections acquired with a tSEM detector. But for our testing data, we acquired HR and LR images of 80 nm sections imaged with a backscatter detector. Based on several metrics including PSNR, SSIM, FRC (Fig. 1), NanoJ-SQUIRREL error mapping analysis (Supplementary Fig. 1)^19^, and visual inspection, we found PSSR very effectively restored the low resolution images (Fig. 1c). Thus, our PSSR model is not restricted to data acquired in the exact same fashion or modality as our training set.

**Fig. 1.**
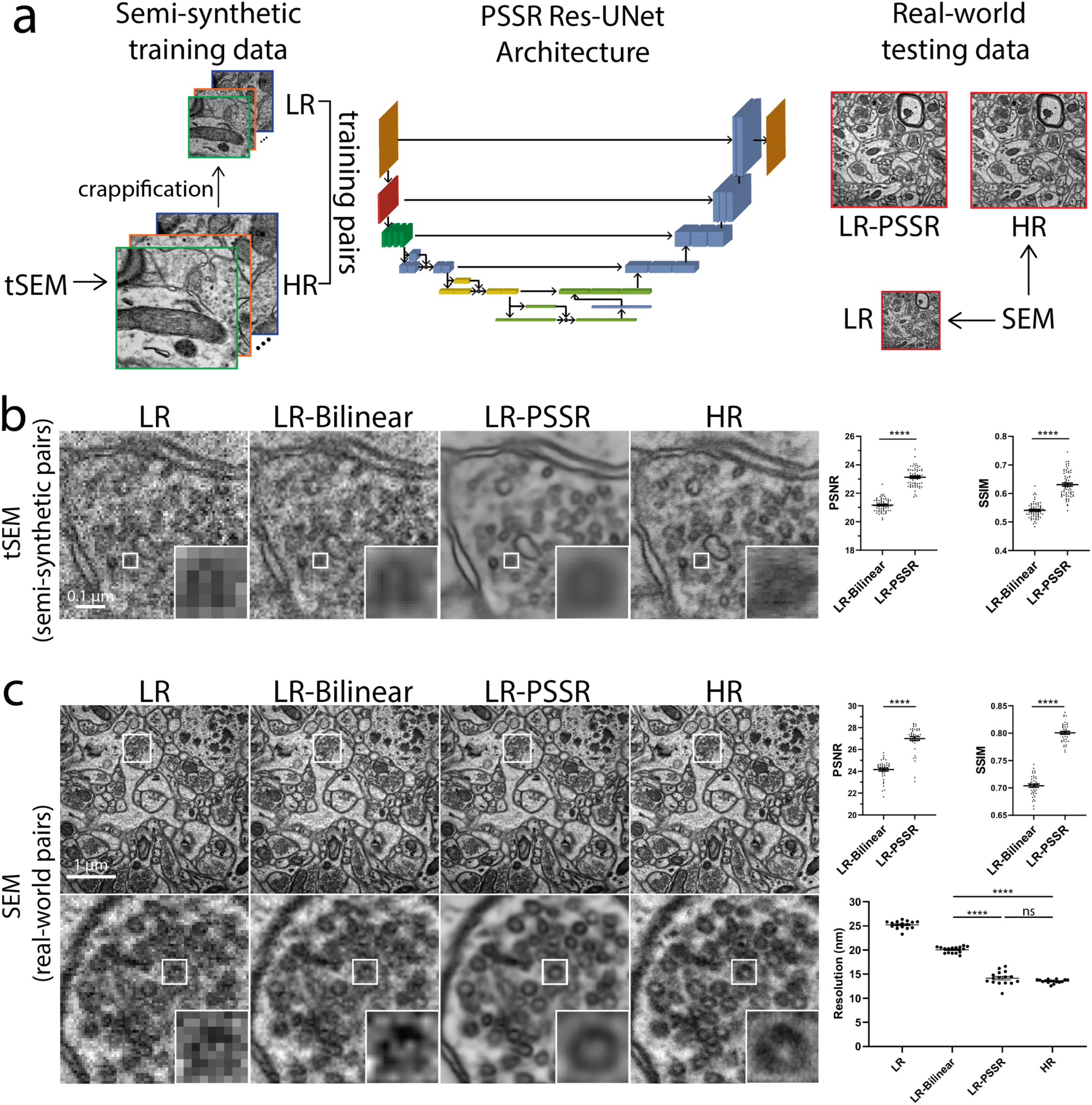
Restoration of semi-synthetic and real-world EM testing data using PSSR model trained on semi-synthetically generated training pairs. **a**, Overview of the general workflow. Training pairs were semi-synthetically created by applying a degrading function to the HR images taken from a scanning electron microscope in transmission mode (tSEM) to generate LR counterparts (left column). Semi-synthetic pairs were used as training data through a dynamic ResNet-based U-Net architecture (middle column). Real-world LR and HR image pairs were both manually acquired under a SEM (right column). The output from PSSR (LR-PSSR) when LR is served as input is then compared to HR to evaluate the performance of our trained model. **b**, Restoration performance on semi-synthetic testing pairs from tSEM. Shown is the same field of view of a representative bouton region from the synthetically created LR input with the pixel size of 8 nm (left column), a 16x bilinear upsampled image with 2 nm pixel size (second column), 16x PSSR upsampled result with 2 nm pixel size (third column) and the HR ground truth acquired at the microscope with the pixel size of 2 nm (fourth column). A close view of the same vesicle in each image is highlighted. The Peak-Signal-to-Noise-Ratio (PSNR) and the Structural Similarity (SSIM) quantification of the semi-synthetic testing sets are shown (right). **c**, Restoration results of manually acquired SEM testing pairs. Shown is the comparison of the LR input acquired at the microscope with a pixel size of 8 nm (left column), 16x bilinear upsampled image (second column), 16x PSSR upsampled output (third column) and the HR ground truth acquired at the microscope with a pixel size of 2 nm (fourth column). Bottom row compares the enlarged region of a presynaptic bouton with one vesicle highlighted in the inset. Graphs comparing PSNR, SSIM and image resolution are also displayed (right). The PSNR and SSIM values were calculated between an upsampled result and its corresponding HR ground truth. Resolution was calculated with the Fourier Ring Correlation (FRC) plugin in NanoJ-SQUIRREL by acquiring two independent images at low and high resolution. All values are shown as mean ± SEM. ns = not significant. *p<0.05, **p<0.01, ***p<0.001, ****p<0.0001; Paired *t*-test.

We next tested whether we could sufficiently restore 8 nm SBFSEM datasets to 2 nm using PSSR, since high quality 2 nm SBFSEM imaging is currently difficult or impossible for us to achieve. Using our PSSR model we were able to restore an 8 nm pixel SBFSEM 3D dataset to 2 nm (Fig. 2a, Supplementary Movie 2). Remarkably, our PSSR model also worked very well on mouse, rat, and fly samples imaged on four different microscopes in four different labs (Fig. 2a-d). In addition to our SBFSEM and SEM imaging systems, PSSR processing appeared to restore images acquired on data from a ZEISS FIB-SEM (from the Hess lab at Janelia Farms, Fig. 2c, Supplementary Movie 3) and a Hitachi Regulus FE-SEM (from the Kubota lab at National Institute for Physiological Sciences). Notably, the PSSR images were much easier to manually segment - a major requirement for properly analyzing 3DEM datasets (Supplementary Movie 4). PSSR processing also performed well on a 10×10×10 nm resolution FIB-SEM fly brain dataset, resulting in a 2 × 2 × 10 nm resolution dataset with higher SNR and resolution (Fig. 2b). Thus, PSSR can be used for 25x super-sampling with useful results, increasing the lateral resolution and speed of FIB-SEM imaging by a factor of at least 25x.

**Fig. 2.**
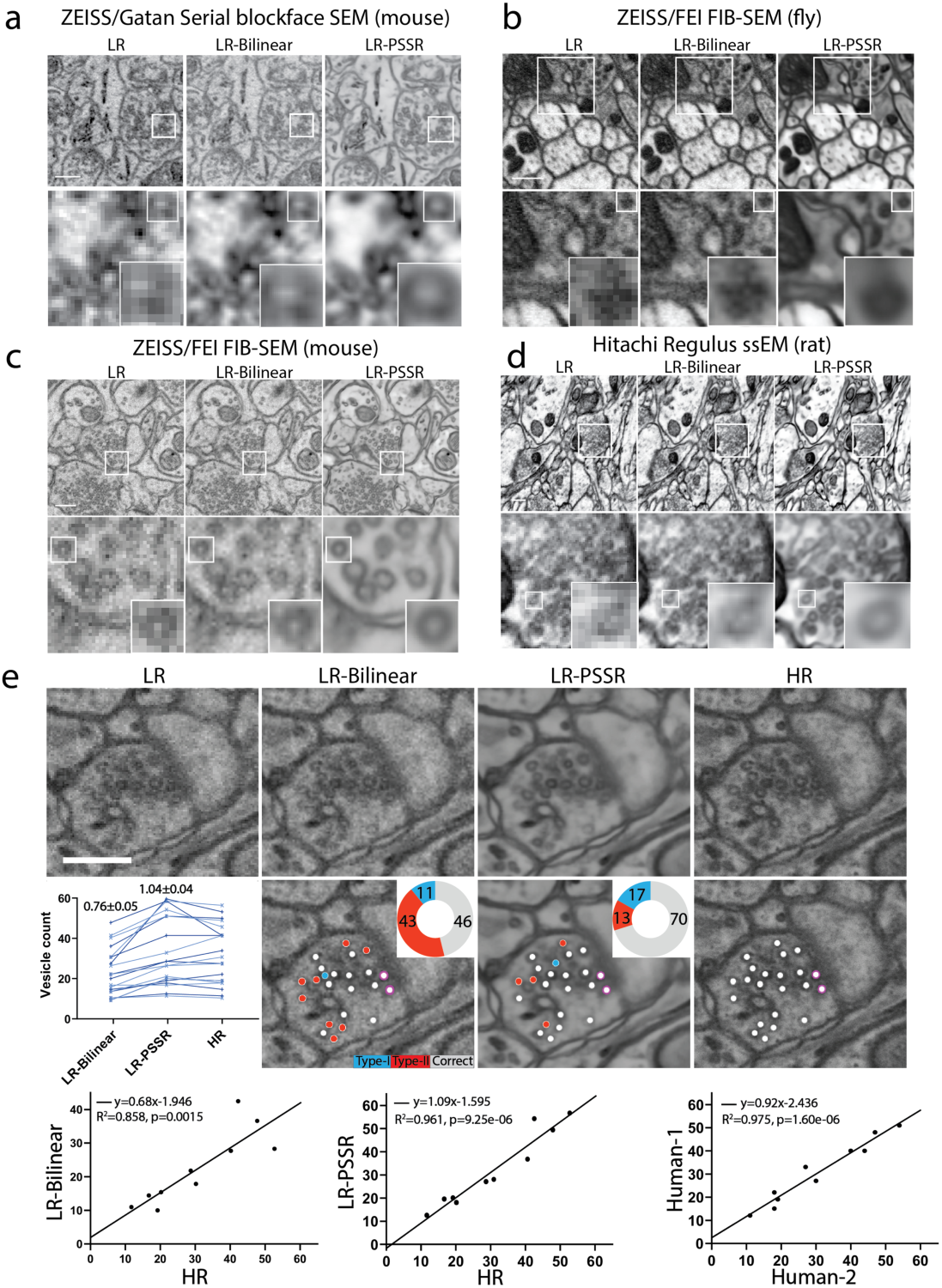
PSSR model is effective for multiple EM modalities and sample types. Shown are representative low resolution (LR), bilinear interpolated (LR-Bilinear) and PSSR-restored (LR-PSSR) images from mouse brain sections imaged with a ZEISS Sigma-VP Gatan Serial Blockface SEM system (**a**), fly sections acquired with ZEISS/FEI focused ion beam-SEM (FIB-SEM) (**b**), mouse sections from ZEISS/FEI FIB-SEM (**c**) and rat sections imaged with a Hitachi Regulus serial section EM (ssEM) (**d**). **e**, Validation of pre-synaptic vesicle detection. LR, LR-Bilinear, LR-PSSR, and ground truth high resolution (HR) images of a representative bouton region as well as their color-labeled vesicle counts are shown. Vesicles colored with red represents false negatives, blue are false positives, and white are true positives. The percentage of each error type is shown in the pie chart. Docked vesicles were labelled with purple dots. A plot of the vesicle counts for two expert humans (light blue and dark blue) for all 3 image sets is shown on the left, with the average total error ± s.e.m. displayed above. The linear regression between LR-Bilinear and HR, LR-PSSR and HR, and two human counters of HR are shown in the third row. The equation for the linear regression, the Goodness-of-Fit (*R*^*2*^) and the *p*-value (*p*) of each graph are displayed. Scale bars = 1.5 μm.

One major concern with deep learning-based image processing is accuracy, and in particular the possibility of false positives (aka “hallucinations”)^4,5,20,21^. As mentioned above, 2 nm pixel SBFSEM datasets are beyond the capabilities for our samples and detector, precluding the generation of ground truth validation images for our SBFSEM data. Given the need to be able to trust processed datasets for which no “ground truth” data exists, we next decided to use ground truth data to determine whether our PSSR output is sufficiently accurate for useful downstream analysis. To do this, we manually acquired low 8 nm and high 2 nm pixel resolution SEM image pairs of ultrathin sections, then 16x super-sampled the 8 nm pixel images (LR) to 2 nm pixel images (HR) using either bilinear interpolation (LR-Bilinear) or PSSR (LR-PSSR). We then measured the PSNR and SSIM of LR-Bilinear and LR-PSSR and found that LR-PSSR significantly outperforms LR-Bilinear. To further test the accuracy and utility of the PSSR output in a more concrete, biological context, we next randomized LR-Bilinear, LR-PSSR, and HR images, then distributed them to two blinded human experts for manual segmentation of presynaptic vesicles, which are both biologically significant and also significantly more difficult to detect with 8nm pixel acquisitions. We found the LR-PSSR segmentation was significantly more accurate than the LR-Bilinear (Fig. 2e). Interestingly, while the LR-PSSR output reduced false negatives by ∼300%, the LR-PSSR output had a slightly higher number of “false positives” than the LR-Bilinear. Most importantly, the variance between the LR-PSSR and HR results was similar to the variance between the two expert human results on HR data (Fig. 2e), which is probably very near maximum possible accuracy and precision. Taken together, our data reveal PSSR to be a viable method for producing 2 nm 3DEM data from 8 nm resolution acquisitions, revealing important subcellular structures that are otherwise lost in many 3DEM datasets. Furthermore, the ability to reliably 16x super-sample lower resolution datasets presents an opportunity to increase the throughput of SEM imaging by at least one order of magnitude.

Similar to SBFSEM, laser scanning confocal microscopy also suffers from a direct relationship between pixel resolution and sample damage (i.e. phototoxicity/photobleaching)^22^. This can be a major barrier for cell biologists who wish to study the dynamics of smaller structures such as mitochondria, which regularly undergo fission and fusion, but also show increased fission and swelling in response to phototoxicity (Supplementary Movie 5, Supplementary Fig. 3). In extreme cases, phototoxicity can cause cell death, which is incompatible with live cell imaging (data not shown). HR scanning confocal microscopy also suffers from the direct relationship between pixel resolution and imaging time, making live cell imaging of faster processes (e.g. organelle motility in neurons) challenging (Supplementary Movie 7). Thus, we sought to determine whether PSSR might provide a viable strategy for increasing the speed and reducing the phototoxicity of live scanning confocal microscopy.

To generate our ground truth training and testing dataset we used a ZEISS Airyscan LSM 880, an advanced confocal microscope that uses a 32-detector array and post-processing pixel reassignment to generate images ∼1.7x higher in resolution (∼120 nm) and ∼8x higher SNR than a standard confocal system. All HR ground truth and training data were acquired with a 63x objective with at least 2x Nyquist pixel size (∼50 ± 10 nm), then Airyscan processed (pixel reassignment) using ZEISS Zen software. Similar to our EM model, semi-synthetic LR training data was generated by computationally degrading HR Airyscan images with random noise and blur. For our LR test data, we acquired images at 16x lower pixel resolution (170 nm) with a 2.5 AU pinhole on a PMT confocal detector, without any additional image processing. We maintained equal pixel dwell time for the HR vs. LR testing acquisitions. We also decreased the laser power for our LR acquisitions by a factor of 4 or 5 (see Supplemental Tables for more details), resulting in a net laser dose decrease of ∼64x to 90x (e.g., 5x lower laser power and 16x shorter exposure time yields a 90x lower laser dose). Furthermore, our LR testing data was not deconvolved and used a much larger effective pinhole size than the HR Airyscan ground truth data. Thus, low resolution, low SNR, undersampled confocal images would need to be restored to oversampled, high SNR, high resolution Airyscan image quality. To start, we trained on live cell timelapses of mitochondria in U2OS cells. As expected, imaging at full resolution (∼49 nm pixels) resulted in significant bleaching and phototoxicity-induced mitochondrial swelling and fission (Supplementary Movie 5). However, the LR (∼196 nm pixels) acquisitions were extremely noisy and pixelated due to undersampling. On the other hand, the LR scans showed far less photobleaching when imaged at the same frame rate and duration (Supplementary Fig. 2). PSSR processing reduced the noise and increased the resolution of the LR acquisitions, as determined by testing on both semi-synthetic and real-world low versus high resolution image pairs (Fig. 3b-c).

**Fig. 3.**
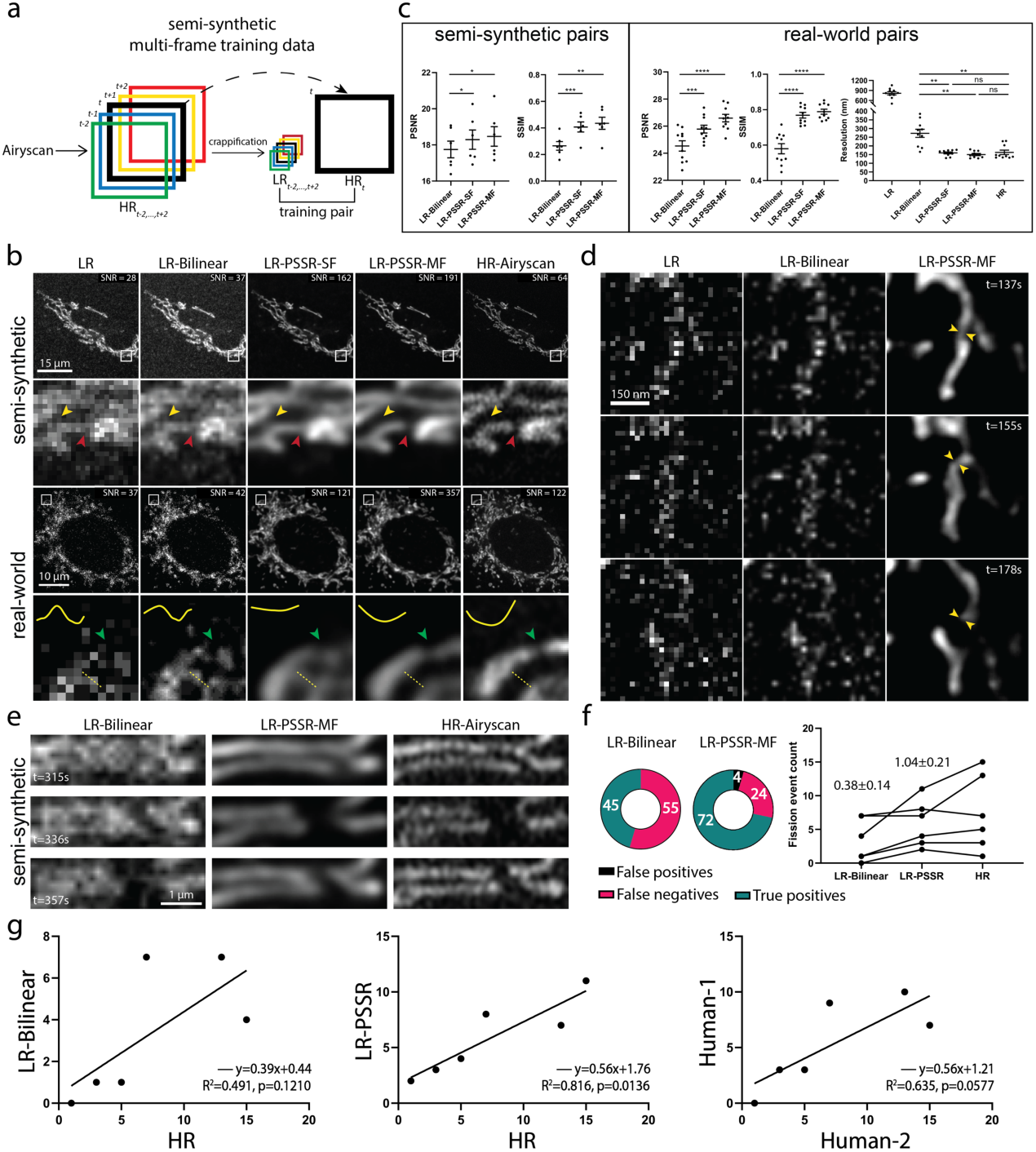
Multi-frame PSSR timelapses of mitochondrial dynamics. **a**, Overview of multi-frame PSSR training data generation method. Five consecutive frames (*HR*_*i*_, *i* ∈ [*t* − 2, *t* +2]) from a HR Airyscan time-lapse movie were synthetically crappified to five LR images (*LR*_*i*_, *i* ∈ [*t* − 2, *t* +2]), which together with the HR middle frame at time *t* (*HR*_*t*_), form a five-to-one training “pair”. **b**, Restoration performance on semi-synthetic and real-world testing pairs. For the semi-synthetic pair, LR was synthetically generated from Airyscan HR movies. Enlarged ROIs show an example of well resolved mitochondrial structures by PSSR, agreeing with Airyscan ground truth images. Multi-frame PSSR (LR-PSSR-MF) further improves the accuracy beyond the standard single-frame PSSR (LR-PSSR-SF) model. Red and yellow arrowheads show two false connecting points in LR-Bilinear and LR-PSSR-SF, which were well separated in LR-PSSR-MF. For the real-world example, ten images of fixed samples were sequentially acquired using a standard confocal (LR) versus an Airyscan (HR) detector with the same pixel dwell time, but 5x lower laser power for the LR acquisition and 16x lower xy pixel resolution, yielding a net reduction in laser dose of ∼90x. The green arrowheads in the enlarged ROIs highlight a well restored gap between two mitochondria segments in the LR-PSSR-MF output. Normalized line-plot cross-section profile (yellow) highlights false bridging between two neighboring structures in LR-Bilinear and LR-PSSR-SF, which was well separated with our PSSR-MF model. Signal-to-Noise Ratio (SNR) measured using the images in both semi-synthetic and real-world examples are indicated. **c**, PSNR and SSIM quantification of the semi-synthetic as well as the real-world testing sets discussed in (b). FRC values measured using two independent low vs. high resolution acquisitions from multiple cells are indicated. **d**, PSSR output captured a transient mitochondrial fission event. Shown is a PSSR-restored dynamic mitochondrial fission event, with three key time frames displayed. Arrows highlight the mitochondrial fission site. **e**, validation of fission event captures using semi-synthetic data. An example of a fission event that was detectable in LR-PSSR but not LR-Bilinear. **f**, For fission event detection, the number of false positives, false negatives, and true positives detected by expert humans was quantified for 6 different timelapses. **g**, The linear regression between LR-Bilinear and HR, LR-PSSR and HR, and two human counters of HR are shown. The equation for the linear regression, the Goodness-of-Fit (R^2^) and the p-value of each graph are displayed. All values are shown as mean ± SEM. ns = not significant. *p<0.05, **p<0.01, ***p<0.001, ****p<0.0001; Paired *t*-test.

For timelapse imaging, one can take advantage of multiframe acquisitions to increase the accuracy of individual frame reconstructions^23,24^. To further improve the performance of PSSR for our timelapse data, we exploited the spatiotemporal continuity of live imaging data and the multi-dimensional capabilities of our PSSR ResU-Net architecture by training on 5 timepoints at a time (MultiFrame-PSSR, or “PSSR-MF”, Fig. 3a). As measured by PSNR, SSIM, and NanoJ-SQUIRREL error mapping, PSSR-MF processing of LR acquisitions (LR-PSSR-MF) showed significantly increased resolution and SNR compared to the raw input (LR), 16x bilinear interpolated input (LR-Bilinear), or single-frame PSSR (LR-PSSR-SF) (Fig. 3b-c, Supplementary Fig. 3). Thus, for all time-lapse PSSR we used PSSR-MF and refer to it as PSSR for the remainder of this article. The improved speed, resolution, and SNR enabled us to detect mitochondrial fission events that were not detectable in the LR or LR-Bilinear images (yellow arrows, Fig. 3d, Supplementary Movie 6). Additionally, the relatively high laser dose during HR acquisitions raises questions as to whether observed fission events are artifacts of phototoxicity. We validated the accuracy of our fission event detection with semi-synthetic data quantified by two expert humans. We found a significant improvement in detecting fission events (56% to 24% false negative rate) with relatively minor increases in false positives (Fig. 3e-g). Thus, PSSR provides an opportunity to detect very fast mitochondrial fission events with fewer phototoxicity-induced artifacts than standard high resolution Airyscan imaging using normal confocal optics and detectors.

In addition to phototoxicity issues, the slow speed of HR scanning confocal imaging results in *<u>temporal</u>* undersampling of fast-moving structures such as motile mitochondria in neurons (Supplementary Fig. 4, Supplementary Movie 8). However, relatively fast LR scans do not provide sufficient pixel resolution or SNR to resolve fission or fusion events, or individual mitochondria when they pass one another along an axon, which can result in faulty analysis or data interpretation (Supplementary Movie 7). Thus, we next tested whether PSSR provided sufficient restoration of undersampled time-lapse imaging of mitochondrial trafficking in neurons.

The overall resolution and SNR improvement provided by PSSR enabled us to resolve adjacent mitochondria moving in an axon, as well as mitochondrial fission and fusion events (Fig. 4a and 4c, Supplementary Fig. 5, Supplementary Movie 8). Since our LR acquisition rates are 16x faster than HR, instantaneous motility details were preserved in LR-PSSR whereas in HR images they were lost (Fig. 4d, Supplementary Fig. 4, Supplementary Movie 8). The overall total distance mitochondria travelled in axons was the same for both LR and HR (Fig. 4f). However, we were able to obtain unique information about how they translocate by imaging at a 16x higher frame rate (Fig. 4g). Interestingly, a larger range of velocities was identified in LR-PSSR than both LR and HR images. Overall, LR-PSSR and HR provided similar values for the percent time mitochondria spent in motion (Fig. 4h). Smaller distances travelled were easier to detect in our LR-PSSR images, and therefore there was an overall reduction in the percent time mitochondria spent in the stopped position in our LR-PSSR data (Fig. 4i). Taken together, these data show PSSR provides a means to detect critical biological events that would not be possible on our confocal system with standard HR or LR imaging.

**Fig. 4.**
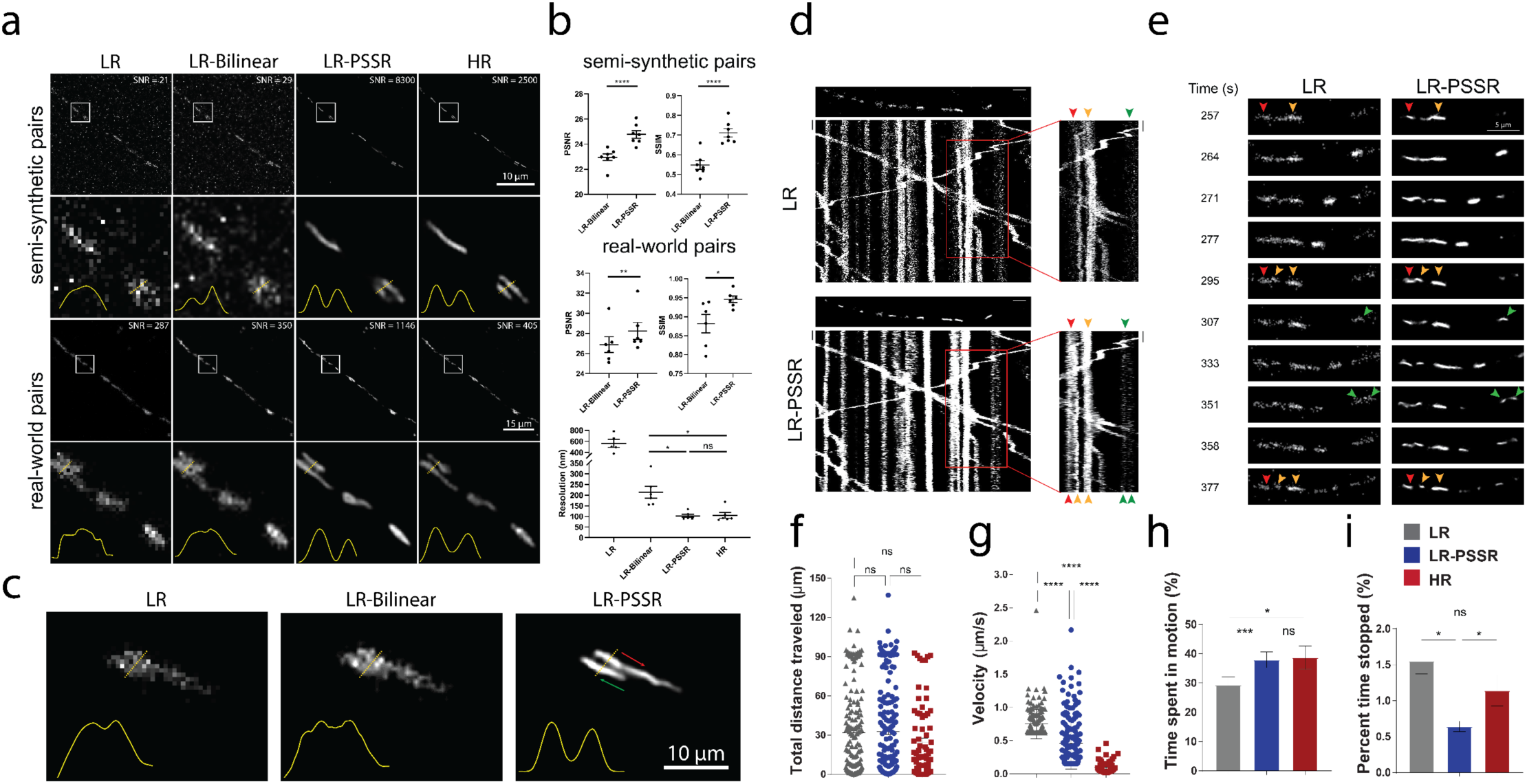
Spatiotemporal analysis of mitochondrial motility in neurons. PSSR facilitates high spatiotemporal resolution imaging of mitochondrial motility in neurons. **a**, Comparison of PSSR results (LR-PSSR) versus bilinear interpolation (LR-Bilinear) on semi-synthetic and real-world testing pairs. Enlarged ROIs from representative images show PSSR resolved two neighboring mitochondria in both semi-synthetic and real-world testing sets, quantified by normalized line plot cross-section profiles. SNR **b**, PSNR (top) and SSIM (middle) quantification of the datasets in (a). FRC resolution measured from two independent acquisitions of the real-world overview dataset discussed in (a) is indicated (bottom). **c**, PSSR restoration of LR timelapses resolves neighboring mitochondria moving past one another in an axon (arrows indicate direction of movement). **d**, Representative kymographs of axonal mitochondrial motility in hippocampal neurons transfected with Mito-dsRed. First frame of each time-lapse movie is shown above each kymograph. Different color arrowheads indicate mitochondria going through fission and fusion events. Each color represents a different mitochondrion. **e**, Enlarged areas of (d), capturing mitochondrial fission and fusion events in real-time. **f-i**, Mitochondrial motility was quantified from time-lapse movies as demonstrated in Supplementary Fig. 4. For each mitochondrial trajectory the total distance mitochondria traveled f, mitochondrial velocity g, percent time mitochondria spent in motion (h) and in pause (i) was quantified. n = 76 - 216 mitochondria from 4 axons and 3 independent transfections. Scale bars = 10 s in d. All values are shown as mean ± SEM. ns = not significant. *p<0.05, **p<0.01, ***p<0.001, ****p<0.0001; Paired *t*- test (b) and Kruskal-Wallis test (f-i).

## Discussion

It is important to consider that any output from a deep learning super-resolution model is a prediction, is never 100% accurate, and is always highly dependent on proper correspondence between the training versus testing data^4,5,21,25^. Whether the level of accuracy of a given model for a given dataset is sufficient is ultimately dependent on the tolerance for error in the measurement being made. For example, our EM PSSR model was validated by segmenting vesicles in presynaptic boutons – the error between our model and the ground truth was similar to the error between two expert humans measuring the ground truth. But we did not rule out the possibility that other structures or regions in the same or similar tissue samples may not be restored by our model with sufficient accuracy. Thus, for each sample type, dataset, and structure of interest, it is essential to first validate the accuracy of the model for the specific task at hand or object(s) of interest before investing in large-scale acquisition or analyses.

That being said, though the accuracy of deep learning approaches such as PSSR is technically lower than “ground truth” data, multiple real-world limitations on acquiring ground truth data may render PSSR the best option. Taken together, our results show the PSSR approach can in principle enable higher speed and resolution imaging with sufficient fidelity for transformative scientific research. All point scanning imaging systems have a direct relationship between pixel resolution and imaging time, sample damage, SNR, and structural detail. Thus, the ability to use deep learning to super-sample under-sampled images provides an opportunity that extends to other point scanning systems, for example ion-based imaging systems^26,27^ or high resolution cryoSTEM^28^.

Interestingly, our semi-synthetic LR images were usually lower quality than our manually acquired LR data. Our real-world LR images were acquired with the same pixel dwell time as our HR data, resulting in 16x lower signal for our real-world LR images. At the same time, our HR-tSEM training data was higher quality than the real-world validation dataset acquired using a backscatter detector on an SEM. This strategy of training a model to attempt to restore lower quality data than needed to a higher quality than possible on a specific testing system appears to be viable for ensuring the model can perform reasonably well on a wide range of real-world data. However, this strategy is still limited by the similarity between the training and testing data. In general, fluorescence data is much sparser and has the potential to be much more variable than EM data. Thus, for fluorescence data it is more important to train and use a specific model for a specific sample type. In all cases, the ability to “crappify” ground truth data to generate semi-synthetic pairs for training greatly increases the throughput for generating useful models. Indeed, any pre-existing high resolution, high SNR dataset of a certain sample type can in theory be used as training data for a deep learning-based image restoration model for that sample type.

It should be noted that we only used MSE loss for our model. For this proof of principle study we wished to generate models that were relatively “naïve” to the particular structures of interest, depending only on pixel-to-pixel information for training and inference. We anticipate employing a more sophisticated loss function could improve the resolution and SNR capabilities of a PSSR model. Using both transfer learning and feature loss presents a practical strategy for optimizing a model for use on a specific dataset or sample type. For future uses of PSSR, we propose an acquisition scheme wherein a relatively limited number of “ground truth” HR images are acquired for fine tuning the model either before or after acquiring the low-resolution experimental data. More importantly - generalized, unsupervised or “self-supervised” denoising approaches^13,29^ as well as deep learning-enabled deconvolution approaches^6,11^ suggest we may one day be able to generate a more generalized model for a specific imaging system, instead of a specific sample type.

Deconvolution methods including structured illumination microscopy, single-molecule localization microscopy, and pixel reassignment microscopy demonstrate the power of configuring optical imaging systems and acquisition schemes with a specific post-processing computational strategy in mind. The power of deep convolutional neural networks for processing image data presents a new opportunity for redesigning imaging systems to exploit these capabilities in order to minimize costs traditionally considered necessary for extracting meaningful imaging data. Similarly, automated, real-time corrections to the images and real-time feedback to the imaging hardware are now easily within reach. This is an area of active investigation in our laboratory and others (Lu Mi, Yaron Meirovitch, Jeff Lichtman, Aravinthan Samuel, Nir Shavit, personal communication).

## Methods

### Semi-synthetic Training Image Generation

HR images were acquired using scanning electron or Airyscan confocal microscopes. Due to the variance of image properties (e.g. format, size, dynamic range and depth) in the acquired HR images, data cleaning is indispensable for generating training sets that can be easily accessed during training. In this article, we differentiate the concept of “data sources” and “data sets”, where data sources refer to uncleaned acquired high resolution images, while data sets refer to images that are generated and preprocessed from data sources. HR data sets were obtained after preprocessing HR images from data sources, LR data sets were generated from HR data sets using a “crappifier” function.

#### Preprocessing

Tiles of predefined sizes (e.g. 256 × 256 and 512 × 512 pixels) were randomly cropped from each frame in image stacks from HR data sources. All tiles were saved as separate images in .*tif* format, which together formed a HR data set.

#### Image Crappification

A “crappifier” was then used to synthetically degrade the HR data sets to LR images, with the goal of approximating the undesired and unavoidable pixel intensity variation in real-world low resolution and low SNR images of the same field of view directly taken under an imaging system. These HR images together with their corrupted counterparts served as training pairs to facilitate “deCrappification”. The crappification function can be simple, but it materially improves both the quality and characteristics of PSSR outputs.

Image sets were normalized from 0 to 1 before being 16x downsampled in pixel resolution (e.g. a 1000 × 1000 pixel image would be downsampled to 250 × 250 pixels). To mimic the image quality degradation caused by 16x undersampling on a real-world point scanning imaging system, salt-and-pepper noise, and Gaussian additive noise with specified local variance were randomly injected into the high-resolution images. The degraded images were then rescaled to 8-bit for viewing with normal image analysis software.

##### EM Crappifier

Random Gaussian-distributed additive noise (*μ*_*EM*_ = 0, *σ*_*EM*_ = 3) was injected. The degraded images were then downsampled using spline interpolation of order 1.

##### MitoTracker and Neuronal Mitochondria Crappifier

The crappification of MitoTracker and neuronal mitochondria data followed a similar procedure. Salt-and-pepper noise was randomly injected in 0.5% of each image’s pixels replacing them with noise, which was followed by the injection of random Gaussian-distributed additive noise (*μ*_*LSM*_ = 0, *σ*_*LSM*_ = 5). The crappified images were then downsampled using spline interpolation of order 1.

#### Data Augmentation

After crappified low-resolution images were generated, we used data augmentation techniques such as random cropping, dihedral affine function, rotation, random zoom to increase the variety and size of our training data^30^.

#### Multi-frame Training Pairs

Unlike imaging data of fixed samples, where we use traditional one-to-one high- and low-resolution images as training pairs, for time-lapse movies, five consecutive frames (*HR*_*i*_, *i* ∈ [*t* − 2, *t* +2]) from a HR Airyscan time-lapse movie were synthetically crappified to five LR images (*LR*_*i*_, *i* ∈ [*t* − 2, *t* +2]), which together with the HR middle frame at time *t* (*HR*_*t*_), form a five-to-one training “pair”. The design of five-to-one training “pairs” leverages the spatiotemporal continuity of dynamic biological behaviors. (Fig. 3a).

### Neural Networks

#### Single-frame Neural Network (PSSR-SF)

A ResNet-based U-Net was used as our convolutional neural network for training^31^. Our U-Net is in the form of encoder-decoder with skip-connections, where the encoder gradually downsizes an input image, followed by the decoder upsampling the image back to its original size. For the EM data, we utilized ResNet pretrained on ImageNet as the encoder. For the design of the decoder, the traditional handcrafted bicubic upscaling filters are replaced with learnable sub-pixel convolutional layers^32^, which can be trained specifically for upsampling each feature map optimized in low-resolution parameter space. This upsampling layer design enables better performance and largely reduces computational complexity, but at the same time causes unignorable checkerboard artifacts due to the periodic time-variant property of multirate upsampling filters^33^. A blurring technique^34^ and a weight initialization method known as “sub-pixel convolution initialized to convolution neural network resize (ICNR)”^35^ designed for the sub-pixel convolution upsampling layers were implemented to remove checkerboard artifacts. In detail, the blurring approach introduces an interpolation kernel of the zero-order hold with the scaling factor after each upsampling layer, the output of which gives out a non-periodic steady-state value, which satisfies a critical condition ensuring a checkerboard artifact-free upsampling scheme^34^. Compared to random initialization, in addition to the benefit of removing checkerboard artifacts, ICNR also empowers the model with higher modeling power and higher accuracy^35^. A self-attention layer inspired by Zhang et al.^36^ was added after each convolutional layer to restore high-frequency details by leveraging larger receptive fields that relate to object shapes. Unlike traditional convolutional neural networks which only use local spatial information to generate high-resolution details, self-attention layers enable global feature extraction to maximize object continuity and consistency.

#### Multi-frame Neural Network (PSSR-MF)

A similar yet slightly modified U-Net was used for time-lapse movie training. The input layer was redesigned to take five frames simultaneously while the last layer still produced one frame as output.

### Training Details

#### Loss Function

MSE loss was used as our loss function.

#### Optimization Methods

Stochastic gradient descent with restarts (SGDR)^37^ was implemented. Aside from the benefits we are able to get through classic stochastic gradient descent, SGDR resets the learning rate to its initial value at the beginning of each training epoch and allows it to decrease again following the shape of a cosine function, yielding lower loss with higher computational efficiency.

#### Cyclic Learning Rate and Momentum

Instead of having a gradually decreasing learning rate as the training converges, we adopted cyclic learning rates^38^, cycling between upper bound and lower bound, which helps oscillate towards a higher learning rate, thus avoiding saddle points in the hyper-dimensional training loss space. In addition, we followed The One Cycle Policy^39^, which restricts the learning rate to only oscillate once between the upper and lower bounds. Specifically, the learning rate linearly increases from the lower bound to the upper bound as the momentum decreases from its upper bound to the lower bound linearly. In the second half of the cycle, the learning rate fits a cosine annealing decreasing from the upper bound to zero while the momentum increases from its lower bound to the upper bound following the same annealing. This training technique achieves superior regularization by preventing the network from overfitting during the middle of the learning process, as well as enables super-convergence^40^ by allowing large learning rates and adaptive momentum.

#### Progressive Resizing (used for EM data only)

Progressive resizing was applied during the training of the EM model. Training was executed in two rounds with HR images scaled to xy pixel sizes of 256 × 256 and 512 × 512 and LR images scaled to 64 × 64 and 128 × 128 progressively. The first round was initiated with an ImageNet pretrained ResU-Net, and the model trained from the first round served as the pre-trained model for the second round. The intuition behind this is it quickly reduces the training loss by allowing the model to see lots of images at a small scale during the early stages of training. As the training progresses, the model focuses more on picking up high-frequency features reflected through fine details that are only stored within larger scale images. Therefore, features that are scale-variant can be recognized through the progressively resized learning at each scale.

#### Discriminative Learning Rates (used for EM data only)

To better preserve the previously learned information, discriminative learning was applied during each round of training for the purpose of fine-tuning. At the first stage of training, only the parameters from the last layer were trainable after loading a pretrained model, which either came from a large-scaled trained publicly available model (i.e., pretrained ImageNet), or from the previous round of training. The learning rate for this stage *lr*_1_ was fixed. Parameters from all layers were set as learnable in the second stage. A linearly spaced learning rate range *lr*_2_ was applied. The learning rate gradually increased across the layers of the entire network architecture. The number of training epochs at each round is noted as [*N*_1_, *N*_2_], where *N*_1_ and *N*_2_ denote the epoch number used at stage one and stage two separately (Supplementary Table 3).

#### Best Model Preservation (used for fluorescence data only)

Instead of saving the last model after training a fixed number of epochs, at the end of each training epoch, PSSR checks if the validation loss goes down compared to the loss from the previous epoch and will only update the best model when a lower loss is found. This technique ensures the best model will never be missed due to local loss fluctuation during the training.

#### Elimination of Tiling Artifacts

Testing images often need to be cropped into smaller tiles before being fed into our model due to the memory limit of graphic cards. This creates tiling edge artifacts around the edges of tiles when stitching them back to the original images. A Gaussian blur kernel (*μ*_*tile*_ = 0, *σ*_*tile*_ = 1) was applied to a 10-pixel wide rectangle region centered in each tiling edge to eliminate the artifacts.

#### Technical specifications

Final models were generated using fast.ai v1.0.55 library (https://github.com/fastai/fastai), PyTorch on two NVIDIA TITAN RTX GPUs. Initial experiments were conducted using NVIDIA Tesla V100s, NVIDIA Quadro p6000s, NVIDIA Quadro M5000s, NVIDIA Titan Vs, NVIDIA GeForce GTX 1080s, or NVIDIA GeForce RTX 2080Ti GPUs.

### Evaluation Metrics

#### PSNR and SSIM quantification

Two classic image quality metrics, PSNR and SSIM, known for their properties of pixel-level data fidelity and perceptual quality fidelity correspondingly, were used for the quantification of our paired testing image sets.

PSNR is inversely correlated with MSE, numerically reflecting the pixel intensity difference between the reconstruction image and the ground truth image, but it is also famous for poor performance when it comes to estimating human perceptual quality. Instead of traditional error summation methods, SSIM is designed to consider distortion factors like luminance distortion, contrast distortion and loss of correlation when interpreting image quality^41^.

#### SNR quantification

SNR was quantified for LSM testing images (Fig. 3b and Fig. 4a) by:

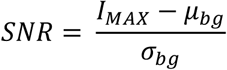

where *I*_*MAX*_ represents the maximum intensity value in the image, *μ*_*bg*_ and *σ*_*bg*_ represent the mean and the standard deviation of the background, respectively^6^.

#### Fourier-Ring-Correlation (FRC) analysis

NanoJ-SQUIRREL^19^ was used to calculate image resolution using FRC method on real-world testing examples with two independent acquisitions of fixed samples (Fig. 1b-c, 3c and Fig. 4b).

#### Resolution Scaled Error (RSE) and Resolution Scaled Pearson’s coefficient (RSP)

NanoJ-SQUIRREL^19^ was used to calculate the RSE and RSP for both semi-synthetic and real-world acquired low (LR), bilinear interpolated (LR-Bilinear), and PSSR (LR-PSSR) images versus ground truth high resolution (HR) images. Difference error maps were also calculated (Supplementary Fig. 1 and 3).

### EM Imaging and Analysis

#### tSEM high resolution training data acquisition

Tissue from a perfused 7-month old Long Evans male rat was cut from the left hemisphere, stratum radiatum of CA1 of the hippocampus. The tissue was stained, embedded, and sectioned at 45 nm using previously described protocols^42^. Sections were imaged using a STEM detector on a ZEISS Supra 40 scanning electron microscope with a 28 kV accelerating voltage and an extractor current of 102 μA (gun aperture 30 μm). Images were acquired with a 2 nm pixel size and a field size of 24576 × 24576 pixels with Zeiss ATLAS. The working distance from the specimen to the final lens was 3.7 mm, and the dwell time was 1.2 μs.

#### EM testing sample preparation and image acquisition

EM data sets were acquired from multiple systems at multiple institutions for this study.

For our testing ground truth data, paired LR and HR images of the adult mouse hippocampal dentate gyrus middle molecular layer neuropil were acquired from ultrathin sections (80 nm) collected on silicon chips and imaged in a ZEISS Sigma VP FE-SEM^15^. All animal work was approved by the Institutional Animal Care and Use Committee (IACUC) of the Salk Institute for Biological Studies. Samples were prepared for EM according the NCMIR protocol^43^. Pairs of 4 × 4 μm images were collected from the same region at pixels sizes of both 8 nm and 2 nm using Fibics ATLAS software (InLens detector; 3 kV; dwell time, 5.3 μs; line averaging, 2; aperture, 30 μm; working distance, 2 mm).

Serial block face scanning electron microscope images were acquired with a Gatan 3View system installed on the ZEISS Sigma VP FE-SEM. Images were acquired using a pixel size of 8 nm on a Gatan backscatter detector at 1 kV and a current of 221 pA. The pixel dwell time was 2 μs with an aperture of 30 μm and a working distance of 6.81 mm. The section thickness was 100 nm and the field of view was 24.5 × 24.5 μm.

Mouse FIB-SEM data sample preparation and image acquisition settings were previously described in the original manuscript the datasets were published^16^. Briefly, the images were acquired with 4 nm voxel resolution. We downsampled the lateral resolution to 8 nm, then applied our PSSR model to the downsampled data to ensure the proper 8-to-2 nm transformation for which the PSSR was trained.

Fly FIB-SEM data sample preparation and image acquisition settings were previously described in the original manuscript the datasets were published^44^. Briefly, images were acquired with 10 nm voxel resolution. We first upsampled the xy resolution to 8 nm using bilinear interpolation, then applied our PSSR model to the upsampled data to ensure the proper 8-to-2 nm transformation for which the PSSR model was originally trained.

The rat SEM data sample was acquired from an 8-week old male Wistar rat that was anesthetized with an overdose of pentobarbital (75 mg kg^-1^) and perfused through the heart with 5 - 10 ml of a solution of 250 mM sucrose 5 mM MgCl_2_ in 0.02 M phosphate buffer (pH 7.4) (PB) followed by 200 ml of 4% paraformaldehyde containing 0.2% picric acid and 1% glutaraldehyde in 0.1 M PB. Brains were then removed and oblique horizontal sections (50 µm thick) of frontal cortex/striatum were cut on a vibrating microtome (Leica VT1200S, Nussloch, Germany) along the line of the rhinal fissure. The tissue was stained and cut to 50 nm sections using ATUMtome (RMC Boeckeler, Tucson, USA) for SEM imaging using the protocol described in the original publication for which the data was acquired^45^. The Hitachi Regulus rat SEM data was acquired using a Regulus 8240 FE-SEM with an acceleration voltage of 1.5 kV, a dwell time of 3 μs, using the backscatter detector with a pixel resolution of 10 × 10 nm. We first upsampled the xy resolution to 8 nm using bilinear interpolation, then applied our PSSR model to the upsampled data to ensure the proper 8-to-2 nm transformation for which the PSSR model was originally trained.

#### EM segmentation and analysis

Image sets generated from the same region of neuropil (LR-Bilinear; LR-PSSR; HR) were aligned rigidly using the ImageJ plugin Linear stack alignment with SIFT^46^. Presynaptic axonal boutons (n = 10) were identified and cropped from the image set. The bouton image sets from the three conditions were then assigned randomly generated filenames and distributed to two blinded human experts for manual segmentation of presynaptic vesicles. Vesicles were identified by having a clear and complete membrane, being round in shape, and of approximately 35 nm in diameter. For consistency between human experts, vesicles that were embedded in or attached to obliquely sectioned axonal membranes were excluded. Docked and non-docked synaptic vesicles were counted as separate pools. Vesicle counts were recorded and unblinded and grouped by condition and by expert counter. Linear regression analyses were conducted between the counts of the HR images and the corresponding images of the two different LR conditions (LR-Bilinear; LR-PSSR) to determine how closely the counts corresponded between the HR and LR conditions. Linear regression analysis was also used to determine the variability between counters.

### Fluorescence Imaging and Analysis

#### U2OS cell culture

U2OS cells were purchased from ATCC. Cells were grown in DMEM supplemented with 10% fetal bovine serum at 37 °C with 5% CO_2_. Cells were plated onto either 8-well #1.5 imaging chambers or #1.5 35 mm dishes (Cellvis) coated with 10 μg/mL fibronectin in PBS at 37 °C for 30 minutes prior to plating. 50 nM MitoTracker Deep Red or CMXRos Red (ThermoFisher) was added for 30 minutes then washed for at least 30 minutes to allow for recovery time before imaging in FluoroBrite (ThermoFisher) media.

#### Airyscan confocal imaging of U2OS cells

Cells were imaged with a 63x 1.4 NA oil objective on a ZEISS 880 LSM Airyscan confocal system with an inverted stage and heated incubation system with 5% CO_2_ control. For both HR and LR images, equal or lower (when indicated) laser power and equal pixel dwell time of ∼1 μs/pixel was used. High resolution Airyscan images (HR) were acquired using 2x Nyquist pixel size of 42 - 59 nm/pixel (depending on the wavelength) in SR mode (i.e. a virtual pinhole size of 0.2 AU), then processed using ZEISS Zen software with auto-filter settings. Low resolution images (LR) were acquired using the same settings but with 0.5x Nyquist pixel size (196 nm/pixel) and a physical pinhole size of 2.5 AU. For testing PSSR-MF, at least 10 sequential frames of fixed samples were acquired with high- and low-resolution settings in order to facilitate PSSR-MF processing.

#### Neuron preparation

Primary hippocampal neurons were prepared from E18 rat (Envigo) embryos as previously described. Briefly, hippocampal tissue was dissected from embryonic brain and further dissociated to single hippocampal neuron by trypsinization with Papain (Roche). The prepared neurons were plated on coverslips (Bellco Glass) coated with 3.33 μg/mL laminin (Life Technologies) and 20 μg/mL poly-L-Lysine (Sigma) at the density of 7.5 × 10^4^ cells/cm^2^. The cells were maintained in Neurobasal medium supplemented with B27 (Life Technologies), penicillin/streptomycin and L-glutamine for 7 - 21 days in vitro. Two days before imaging, the hippocampal neurons were transfected with Lipofectamine 2000 (Life Technologies).

#### Fission event detection and analysis

Given fission events cannot be precisely defined across different evaluators, a HR timelapse of mitotracker stained cells was first given to two human experts as a pilot experiment to examine and correct the inspection performance of all experts. Three conditions (LR-Bilinear; LR-PSSR; HR) of the same Airyscan timelapses (n = 6) were then sequentially assigned to two blinded human experts for mitochondria fission event detection. Fission event counts were recorded and unblinded and grouped by condition and by expert counter. Linear regression analyses were conducted between the counts of the HR images and the corresponding images of the two different LR conditions (LR-Bilinear; LR-PSSR) to determine how closely the counts corresponded between the HR and LR conditions. Linear regression analysis was used to determine the variability between counters.

#### Neuronal mitochondria imaging and kymograph analysis

Live-cell imaging of primary neurons was performed using a Zeiss LSM 880 confocal microscope, enclosed in a temperature control chamber at 37 °C and 5% CO_2_, using a 63x (NA 1.4) oil objective in SR-Airyscan mode (i.e. 0.2 AU virtual pinhole). For low resolution conditions, images were acquired with a confocal PMT detector with a pinhole size of 2.5 AU at 440 × 440 pixels at 0.5x Nyquist (170 nm/pixel) every 270.49 ms using a pixel dwell time of 1.2 µs and a laser power ranging between 1 - 20 µW. For high resolution conditions, images were acquired at 1764 × 1764 pixels at 2x Nyquist (∼42 nm/pixel) every 4.33 s using a pixel dwell time of 1.2 µs and a laser power of 20 µW. Imaging data were collected using Zen Black software. High resolution images were further processed using Zen Blue’s 2D-SR Airyscan processing. Time-lapse movies were analyzed by a custom-written ImageJ macro Kymolyzer as described previously^47^.

#### Fluorescence photobleaching quantification

Normalized mean intensity over time was measured using Fiji software. Given a time-lapse image stack with *N* slices, a background region was randomly selected and remained unchanged across frames. The normalized mean intensity 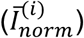 can be expressed as:

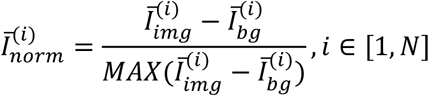

where *i* represents the frame index, 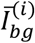 represents the mean intensity of the selected background region at frame *i* and 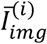 represents the intensity mean of the entire frame *i*.

## Acknowledgements

The authors thank John Sedat, Terry Sejnowski, Antonio Pinto-Duarte, Florian Jug, Martin Weigert, Kirti Prakash, Stephan Saalfeld, and the entire NSF NeuroNex consortium for invaluable advice and critical feedback on our data and the manuscript. The authors also thank Harald Hess and Shan Xu for sharing their FIB-SEM data. U.M., L.F., T.Z., and S.W.N. are supported by the Waitt Foundation and NIH-NCI P30 Grant No. 014195. J.H. and F.M. are supported by the Wicklow AI in Medicine Research Initiative. K.M.H. is supported by NSF Grant No. 1707356 and NIH/NIMH Grant No. 2R56MH095980-06. Research in the laboratory of G.P. is supported by the University of California San Diego institutional funds, Parkinson’s Foundation (PF-JFA-1888), and NIH Grant No. R35GM128823. S.B.Y. is funded by NIH Grant No. T32GM007240. Y.K. was supported by Japan Society for the Promotion of Science KAKENHI Grant 17H06311. The authors acknowledge the Texas Advanced Computing Center (TACC) at The University of Texas at Austin for providing GPU resources that have contributed to the research results reported within this paper. We also gratefully acknowledge the support of NVIDIA Corporation with the donation of the NVIDIA Quadro M5000 and NVIDIA Titan V used for this research.

## Additional information

### Code availability

PSSR source code and documentation are available for download on GitHub (https://github.com/BPHO-Salk/PSSR) and are free for non-profit use.

### Data availability

Example training data and pretrained models are included in the GitHub release (https://github.com/BPHO-Salk/PSSR). In the near future, the entirety of our training and testing data sets and data sources will be made available via a publicly available image hosting website.

## Supplementary Figures

**Supplementary Fig. 1.**
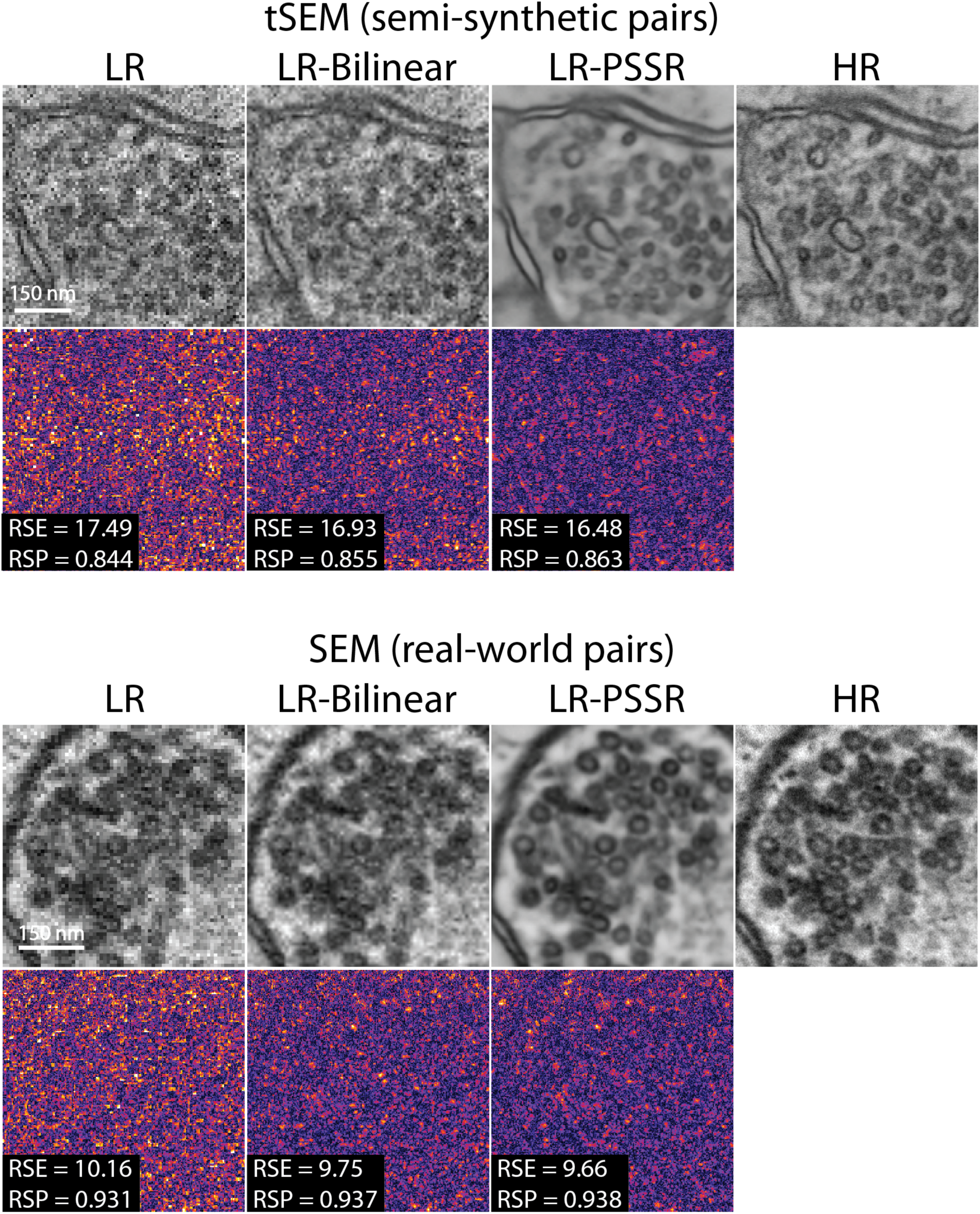
NanoJ-SQUIRREL error-maps of PSSR images. NanoJ-SQUIRREL was used to calculate the resolution scaled error (RSE) and resolution scaled Pearson’s coefficient (RSP) for both semi-synthetic and real-world acquired low (LR), bilinear interpolated (LR-Bilinear), and PSSR (LR-PSSR) images versus ground truth high resolution (HR) images. For these representative images from Fig. 1, the RSE and RSP images are shown along with the difference images for each output.

**Supplementary Fig. 2.**
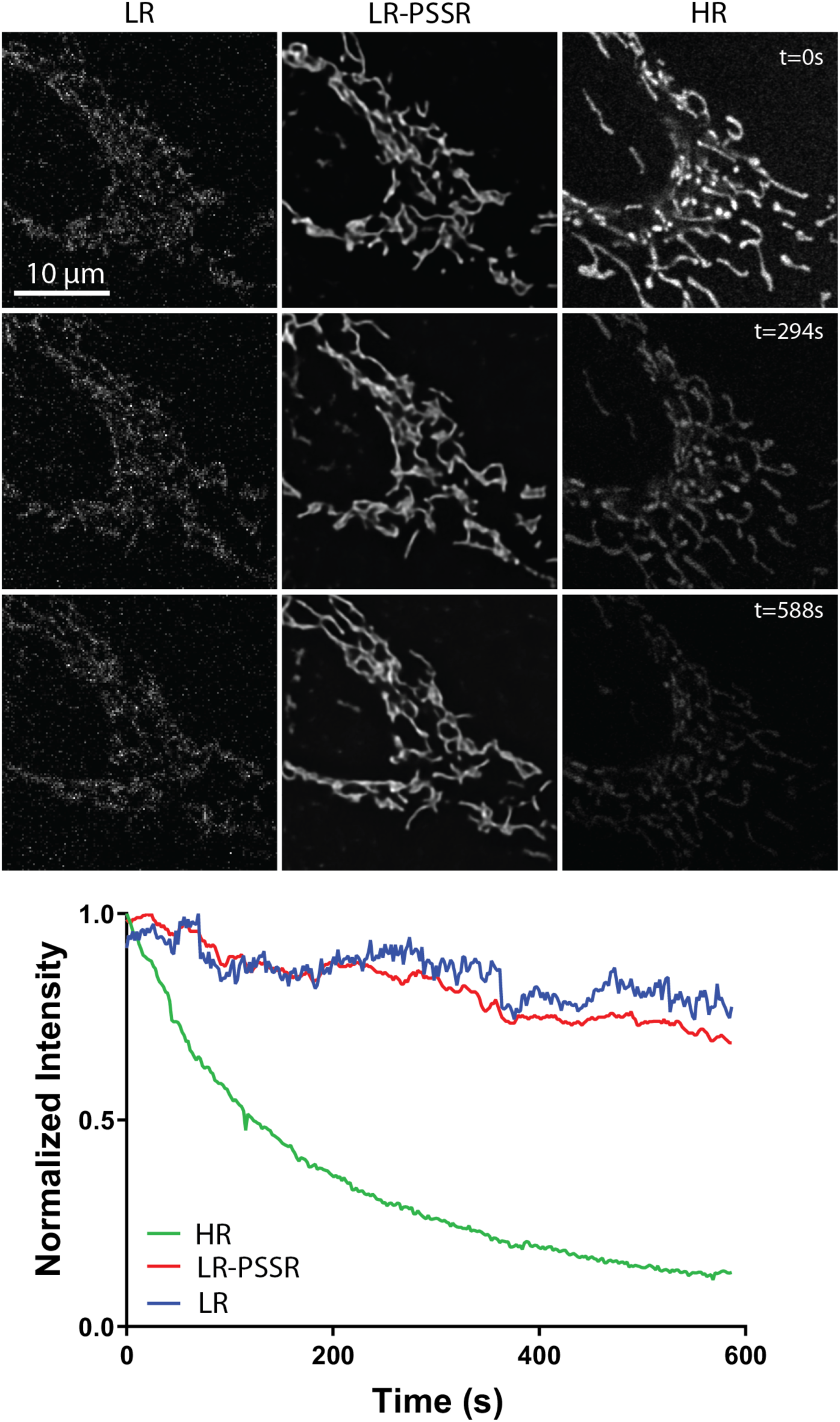
Undersampling significantly reduces photobleaching. U2OS cells stained with mitotracker were imaged every 2 seconds with the same laser power (2.5 μW) and pixel dwell time (∼1 μs), but with 16x lower resolution (196 × 196 nm xy pixel size) than full resolution Airyscan acquisitions (∼49 × 49 nm xy pixel size). Mean intensity plots show the relative rates of fluorescence intensity loss over time (i.e. photobleaching) for LR, LR-PSSR, and HR images.

**Supplementary Fig. 3.**
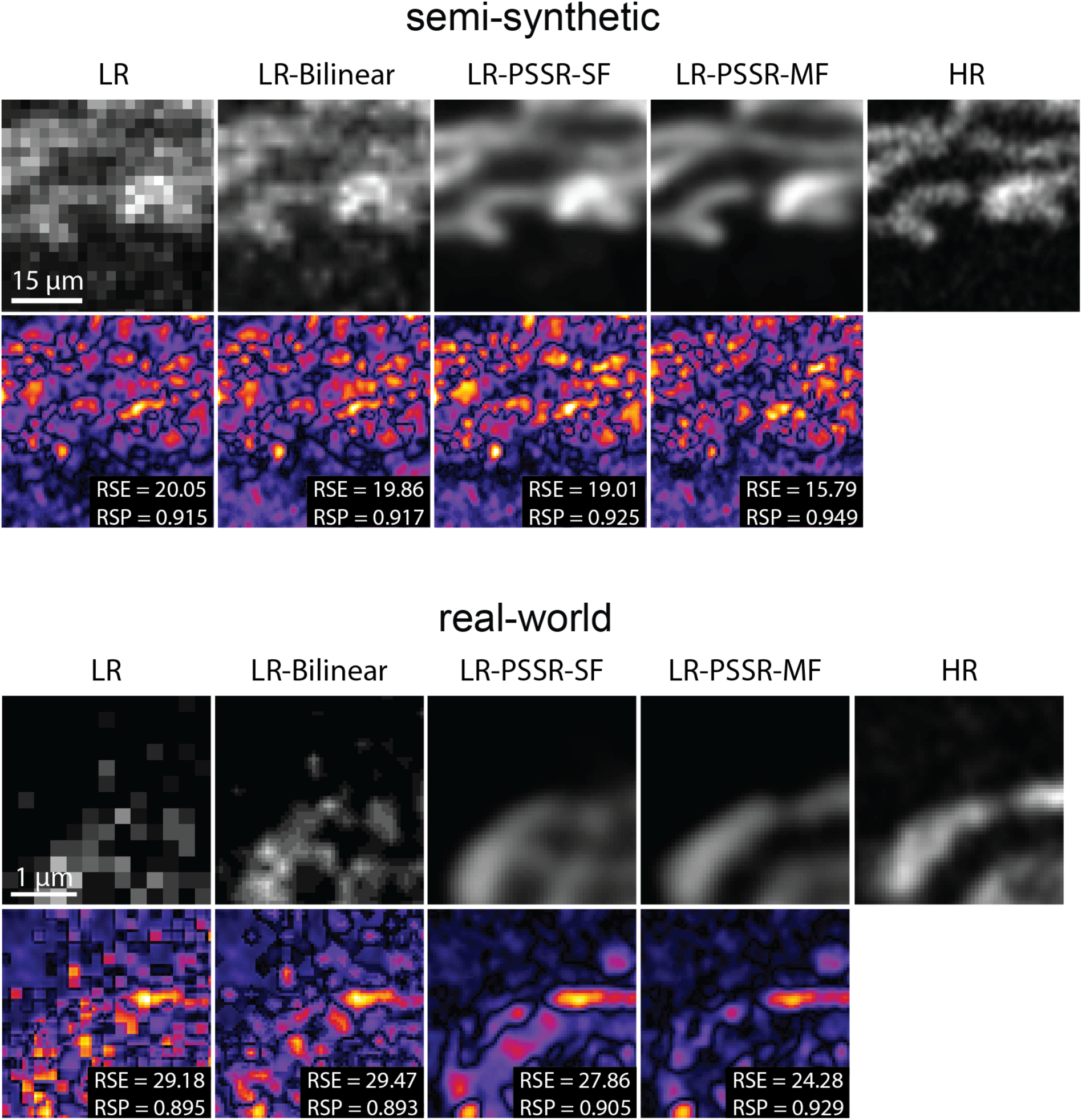
NanoJ-SQUIRREL error-maps of PSSR images. NanoJ-SQUIRREL was used to calculate the resolution scaled error (RSE) and resolution scaled Pearson’s coefficient (RSP) for both semi-synthetic and real-world acquired low (LR), bilinear interpolated (LR-Bilinear), and PSSR (LR-PSSR) images versus ground truth high resolution (HR) images. For these representative images from Fig. 3, the RSE and RSP images are shown along with the difference images for each output.

**Supplementary Fig. 4.**
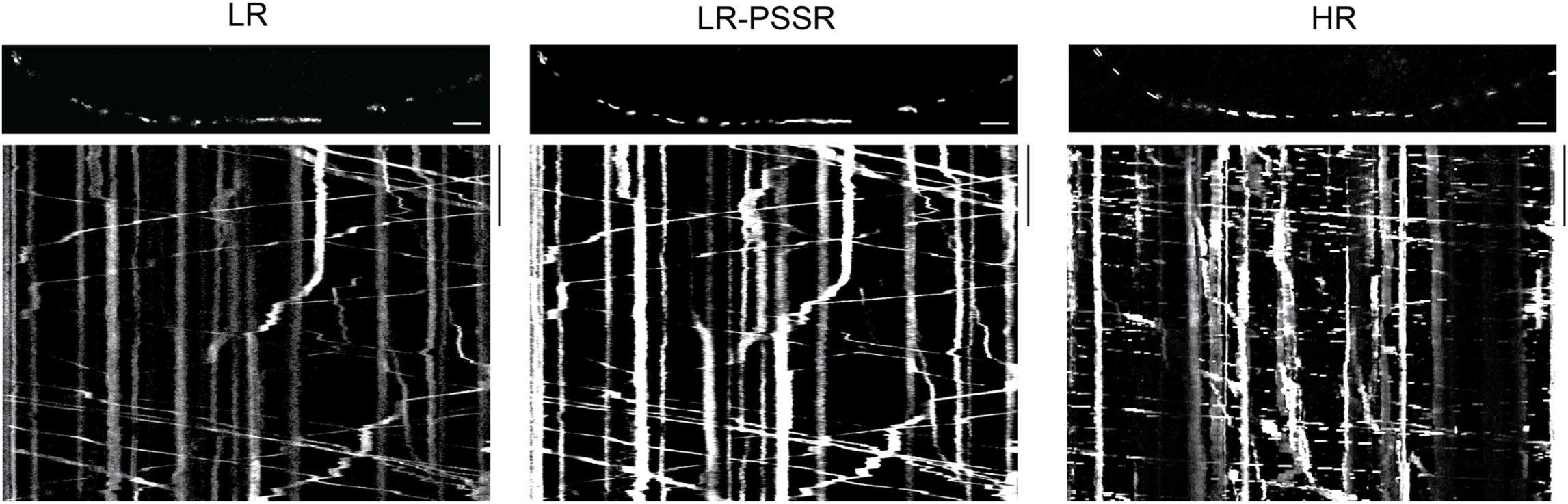
Kymographs of LR vs HR acquisitions. Kymographs generated from timelapses acquired using the same pixel dwell time at low (LR and LR-PSSR) versus high resolution (HR). Note that the HR acquisition was acquired at a different time than the LR acquisitions, and thus the mitochondria are in different positions. The kymograph of the mitochondria in the HR acquisition highlights temporal undersampling as evidenced by the dotted pattern of mitochondrial positions over time, in contrast to LR acquisitions wherein mitochondrial tracks are continuous. Scale bars = 5 μm and 10 s.

**Supplementary Fig. 5.**
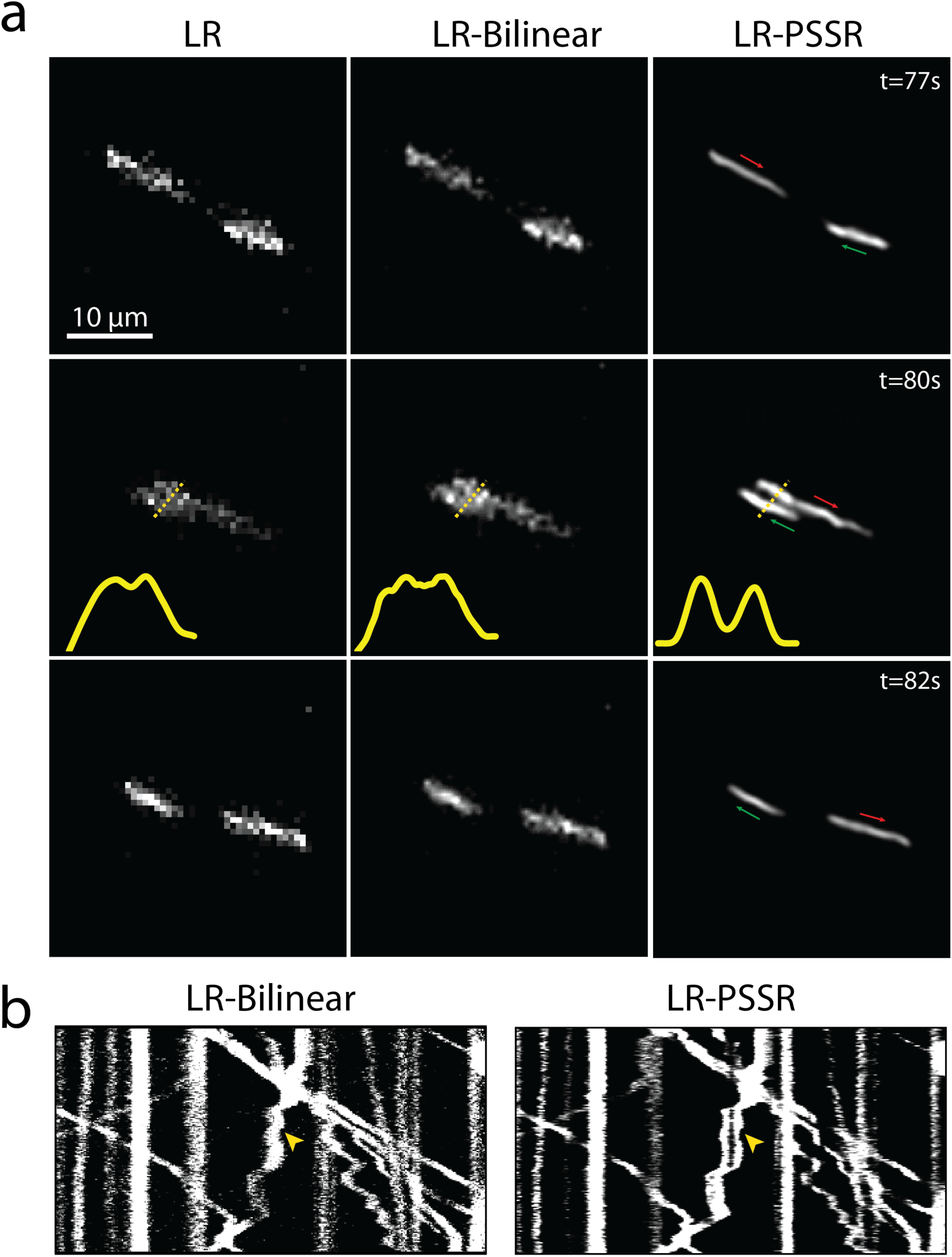
PSSR facilitates detection of mitochondrial motility and dynamics. Rat hippocampal neurons expressing mito-dsRed were undersampled with a confocal detector using 170 nm pixel resolution (LR) to facilitate faster frame rates, then restored with PSSR (LR-PSSR). **a**, before and after time points of the event shown in main Fig. 4 wherein two adjacent mitochondria pass one another but cannot be resolved in the original low resolution (LR) or bilinear interpolated (LR-Bilinear) image but are clearly resolved in the LR-PSSR image. **b**, kymographs of a LR vs LR-PSSR timelapse that facilitates the detection of a mitochondrial fission event (yellow arrow).

**Supplementary Movie 1.** Comparison of high- and low-resolution serial blockface SEM (SBFSEM) 3View acquisition with 2 nm and 8 nm pixel resolutions. In the 2 nm pixel size image stack, high contrast enabled by relatively higher electron doses ensured high resolution and high SNR, which unfortunately at the same time caused severe sample damage, resulting in a failure to serially section the tissue after imaging the blockface. On the other hand, low resolution acquisition at 8 nm pixel size facilitated serial blockface imaging, but the compromised resolution and SNR made it impossible to uncover finer structures in the sample.

**Supplementary Movie 2.** Image restoration achieved by a tSEM-trained PSSR model enables higher resolution SBFSEM imaging. Shown are the lower resolution SBFSEM acquisition input (left) and the PSSR output (right).

**Supplementary Movie 3.** Resolution restoration achieved by tSEM-trained PSSR model enables higher resolution FIB-SEM acquisition. Shown are the lower resolution FIB-SEM acquisition input (left) and the PSSR output (right).

**Supplementary Movie 4.** PSSR facilitates efficient 3D segmentation and reconstruction. Shown is the rendering of the 3D reconstruction of multiple biological structures using the PSSR processed FIB-SEM stack shown in Fig. 2 and Supplementary Movie 3. Specifically, this reconstruction includes mitochondria (purple), endoplasmic reticulum (yellow), presynaptic vesicles (gray), the postsynaptic neuron’s plasma membrane (blue), the postsynaptic density (red) and the presynaptic neuron’s plasma membrane (green). Segmentation was implemented in Reconstruct^48^. Mesh was generated with CellBlender^49,50^. Overlay of the image stack was done using Neuromorph^51^. The animation was made using Blender^52^.

**Supplementary Movie 5.** Photobleaching and cell stress due to high laser dose during high-resolution live cell imaging. Shown is a 10-minute high-resolution time-lapse movie of a U2OS cell stained with Mitotracker Red imaged with an Airyscan microscope. The live-cell acquisition suffered from photobleaching and phototoxicity as reflected by the steadily decreasing fluorescence intensity over time as well as the swelling and fragmenting mitochondria. Imaging condition: 35 μW laser power, 2 s frame rate, 1.15 μm pixel size.

**Supplementary Movie 6.** PSSR restores resolution and SNR to Airyscan equivalent quality with no bleaching and higher imaging speed. Shown are PSSR restoration output (right, ∼49 nm pixels) and its comparison to low resolution acquisition input (left, ∼196 nm pixels). The digitally magnified region highlights a mitochondrial fission event much more easily detected in the PSSR output.

**Supplementary Movie 7.** Comparison of high resolution Airyscan and low resolution confocal time-lapse acquisition of neuronal mitochondria. Corresponding kymographs are also displayed to illustrate the difference in temporal resolution. The Airyscan acquisition has higher spatial resolution but lower temporal resolution due to lower imaging speed, while confocal acquisition gives higher temporal resolution but lower spatial resolution.

**Supplementary Movie 8.** Comparison of PSSR (right) versus bilinear interpolation (left). The enlarged region highlights two adjacent mitochondria passing one another in an axon, the process of which was only resolved in PSSR. Line plot shows the normalized fluorescence intensity of the indicated cross-section.

## Supplementary Tables

**Supplementary Table 1:** Details of fluorescence PSSR training experiments.

| Experiment | U2OS MitoTracker<br>PSSR-SF | U2OS MitoTracker<br>PSSR-MF | Neuron Mito-dsRed<br>PSSR-MF |
| --- | --- | --- | --- |
| Input size (x, y, z) | (128, 128, 1) | (128, 128, 5) | (128, 128, 5) |
| Output size (x, y, z) | (512, 512, 1) | (512, 512, 1) | (512, 512, 1) |
| No. of image pairs for training | 5000 | 5000 | 3000 |
| No. of image pairs for validation | 200 | 200 | 300 |
| No. of GPUs | 2 | 2 | 2 |
| Training datasource size (GB) | 9.4 | 9.4 | 9.5 |
| Validation datasource size (GB) | 0.44 | 0.44 | 1.8 |
| Training dataset size (GB) | 1.38 | 1.7 | 1.03 |
| Validation dataset size (GB) | 0.06 | 0.07 | 0.1 |
| Batch size per GPU | 8 | 8 | 6 |
| No. of epochs | 100 | 100 | 50 |
| Training time (h) | 3.76 | 3.77 | 1.25 |
| Learning rate | 4e-4 | 4e-4 | 4e-4 |
| Best model found at epoch | 33 | 86 | 39 |
| Normalized to ImageNet statistics? | No | No | No |
| ResNet size | ResNet34 | ResNet34 | ResNet34 |
| Loss function | MSE | MSE | MSE |

**Supplementary Table 2:** Details of fluorescence PSSR testing data for PSNR, SSIM, and error-mapping.

|  | U2OS MitoTracker |  |  | Neuron Mito-dsRed |  |  |
| --- | --- | --- | --- | --- | --- | --- |
|  | Semi-synthetic | Real-world |  | Semi-synthetic | Real-world |  |
|  | HR | LR | HR | HR | LR | HR |
| Microscopy | Airyscan | Confocal | Airyscan | Airyscan | Confocal | Airyscan |
| Laser power (μW) | 35 | 7 | 28 | 82 | 11 | 55 |
| Data source size (MB) | 1250 | 7.15 | 200 | 5920 | 10.7 | 305 |
| Data set size (MB) | 325 | 6.39 | 191 | 1970 | 10.4 | 305 |
| Total number of different cells | 6 | 10 | 10 | 7 | 10 | 10 |

**Supplementary Table 3:** Details of the EM PSSR training experiment.

| EM training data |  |  |  |
| --- | --- | --- | --- |
| Round of progressive resizing | 1 | 2 | 3 |
| Input size ($x, y, z$) | (32, 32, 3) | (64, 64, 3) | (128, 128, 3) |
| Output size ($x, y, z$) | (128,128,3) | (256, 256, 3) | (512, 512, 3) |
| Batch size per GPU | 64 | 16 | 8 |
| No. of epochs [$N_1, N_2$] | [1,1] | [3,3] | [3,3] |
| Learning rate [$lr_1, lr_2$] | [1e-3, (1e-5,1e-3)] | [1e-3, (1e-5,1e-3)] | [1e-3, (1e-5,1e-4)] |
| Training dataset size (GB) | 3.9 | 15.56 | 62.24 |
| Validation dataset size (GB) | 0.96 | 3.89 | 15.56 |
| Training datasource size (GB) | 105 |  |  |
| Validation datasource size (GB) | 26 |  |  |
| No. of image pairs for training | 80,000 |  |  |
| No. of image pairs for validation | 20,000 |  |  |
| Total training time (h) | 16 |  |  |
| Normalized to ImageNet statistics? | Yes |  |  |
| ResNet size | ResNet34 |  |  |
| Loss function | MSE |  |  |
| No. of GPUs | 2 |  |  |

## References

1 Wang, Z., Chen, J. & Hoi, S. C. H. Deep Learning for Image Super-resolution: A Survey. eprint 1902.06068, arXiv:1902.06068 (2019).

2 Jain, V. et al. Supervised Learning of Image Restoration with Convolutional Networks. (2007).

3 Romano, Y., Isidoro, J. & Milanfar, P. RAISR: rapid and accurate image super resolution. IEEE Transactions on Computational Imaging 3, 110–125 (2016).

4 Moen, E. et al. Deep learning for cellular image analysis. Nat Methods, doi:10.1038/s41592-019-0403-1 (2019).

5 Weigert, M. et al. Content-aware image restoration: pushing the limits of fluorescence microscopy. Nat Methods 15, 1090–1097, doi:10.1038/s41592-018-0216-7 (2018).

6 Wang, H. et al. Deep learning enables cross-modality super-resolution in fluorescence microscopy. Nat Methods 16, 103–110, doi:10.1038/s41592-018-0239-0 (2019).

7 Ouyang, W., Aristov, A., Lelek, M., Hao, X. & Zimmer, C. Deep learning massively accelerates super-resolution localization microscopy. Nat Biotechnol 36, 460–468, doi:10.1038/nbt.4106 (2018).

8 Nelson, A. J. & Hess, S. T. Molecular imaging with neural training of identification algorithm (neural network localization identification). Microsc Res Tech 81, 966–972, doi:10.1002/jemt.23059 (2018).

9 Li, Y. et al. DLBI: deep learning guided Bayesian inference for structure reconstruction of super-resolution fluorescence microscopy. Bioinformatics 34, i284–i294, doi:10.1093/bioinformatics/bty241 (2018).

10 Buchholz, T. O. et al. Content-aware image restoration for electron microscopy. Methods Cell Biol 152, 277–289, doi:10.1016/bs.mcb.2019.05.001 (2019).

11 Guo, M. et al. Accelerating iterative deconvolution and multiview fusion by orders of magnitude. bioRxiv, 647370, doi:10.1101/647370 (2019).

12 Buchholz, T.-O., Jordan, M., Pigino, G. & Jug, F. in 2019 IEEE 16th International Symposium on Biomedical Imaging (ISBI 2019). 502-506 (IEEE).

13 Batson, J. & Royer, L. Noise2self: Blind denoising by self-supervision. arXiv preprint 1901.11365 (2019).

14 Heinrich, L., Bogovic, J. A. & Saalfeld, S. in Medical Image Computing and Computer-Assisted Intervention - MICCAI 2017. (eds Maxime Descoteaux et al.) 135–143 (Springer International Publishing).

15 Horstmann, H., Korber, C., Satzler, K., Aydin, D. & Kuner, T. Serial section scanning electron microscopy (S3EM) on silicon wafers for ultra-structural volume imaging of cells and tissues. PLoS One 7, e35172, doi:10.1371/journal.pone.0035172 (2012).

16 Xu, C. S. et al. Enhanced FIB-SEM systems for large-volume 3D imaging. Elife 6, doi:10.7554/eLife.25916 (2017).

17 Denk, W. & Horstmann, H. Serial block-face scanning electron microscopy to reconstruct three-dimensional tissue nanostructure. PLoS Biol 2, e329, doi:10.1371/journal.pbio.0020329 (2004).

18 Kuwajima, M., Mendenhall, J. M., Lindsey, L. F. & Harris, K. M. Automated transmission-mode scanning electron microscopy (tSEM) for large volume analysis at nanoscale resolution. PLoS One 8, e59573, doi:10.1371/journal.pone.0059573 (2013).

19 Culley, S. et al. Quantitative mapping and minimization of super-resolution optical imaging artifacts. Nat Methods 15, 263–266, doi:10.1038/nmeth.4605 (2018).

20 von Chamier, L., Laine, R. F. & Henriques, R. Artificial intelligence for microscopy: what you should know. Biochem Soc Trans, doi:10.1042/BST20180391 (2019).

21 Belthangady, C. & Royer, L. A. Applications, promises, and pitfalls of deep learning for fluorescence image reconstruction. Nat Methods, doi:10.1038/s41592-019-0458-z (2019).

22 Phototoxicity revisited. Nat Methods 15, 751, doi:10.1038/s41592-018-0170-4 (2018).

23 Wronski, B. et al. Handheld Multi-Frame Super-Resolution. arXiv preprint 1905.03277 (2019).

24 Huang, X. et al. Fast, long-term, super-resolution imaging with Hessian structured illumination microscopy. Nat Biotechnol 36, 451–459, doi:10.1038/nbt.4115 (2018).

25 Barbastathis, G., Ozcan, A. & Situ, G. On the use of deep learning for computational imaging. Optica 6, 921–943, doi:10.1364/OPTICA.6.000921 (2019).

26 Keren, L. et al. MIBI-TOF: A multiplexed imaging platform relates cellular phenotypes and tissue structure. Sci Adv 5, eaax5851, doi:10.1126/sciadv.aax5851 (2019).

27 Arrojo, E. D. R. et al. Age Mosaicism across Multiple Scales in Adult Tissues. Cell Metab 30, 343–351 e343, doi:10.1016/j.cmet.2019.05.010 (2019).

28 Wolf, S. G. & Elbaum, M. CryoSTEM tomography in biology. Methods Cell Biol 152, 197–215, doi:10.1016/bs.mcb.2019.04.001 (2019).

29 Krull, A., Buchholz, T.-O. & Jug, F. in Proceedings of the IEEE Conference on Computer Vision and Pattern Recognition. 2129–2137.

30 Perez, L. & Wang, J. The effectiveness of data augmentation in image classification using deep learning. arXiv preprint 1712.04621 (2017).

31 Ronneberger, O., Fischer, P. & Brox, T. in International Conference on Medical image computing and computer-assisted intervention. 234–241 (Springer).

32 Shi, W. et al. in Proceedings of the IEEE conference on computer vision and pattern recognition. 1874–1883.

33 Harada, Y., Muramatsu, S. & Kiya, H. in 9th European Signal Processing Conference (EUSIPCO 1998). 1–4 (IEEE).

34 Sugawara, Y., Shiota, S. & Kiya, H. in 2018 25th IEEE International Conference on Image Processing (ICIP). 66–70 (IEEE).

35 Aitken, A. et al. Checkerboard artifact free sub-pixel convolution: A note on sub-pixel convolution, resize convolution and convolution resize. arXiv preprint 1707.02937 (2017).

36 Zhang, H., Goodfellow, I., Metaxas, D. & Odena, A. Self-attention generative adversarial networks. arXiv preprint 1805.08318 (2018).

37 Loshchilov, I. & Hutter, F. Sgdr: Stochastic gradient descent with warm restarts. arXiv preprint 1608.03983 (2016).

38 Smith, L. N. in 2017 IEEE Winter Conference on Applications of Computer Vision (WACV). 464–472 (IEEE).

39 Smith, L. N. A disciplined approach to neural network hyper-parameters: Part 1--learning rate, batch size, momentum, and weight decay. arXiv preprint 1803.09820 (2018).

40 Smith, L. N. & Topin, N. Super-convergence: Very fast training of residual networks using large learning rates. (2018).

41 Hore, A. & Ziou, D. in 2010 20th International Conference on Pattern Recognition. 2366–2369 (IEEE).

42 Kuwajima, M., Mendenhall, J. M. & Harris, K. M. Large-volume reconstruction of brain tissue from high-resolution serial section images acquired by SEM-based scanning transmission electron microscopy. Methods Mol Biol 950, 253–273, doi:10.1007/978-1-62703-137-0_15 (2013).

43 Deerinck, T. J., Bushong, E. A., Thor, A. & Ellisman, M. H. NCMIR methods for 3D EM: a new protocol for preparation of biological specimens for serial block face scanning electron microscopy. Microscopy, 6–8 (2010).

44 Takemura, S. Y. et al. Synaptic circuits and their variations within different columns in the visual system of Drosophila. Proc Natl Acad Sci U S A 112, 13711–13716, doi:10.1073/pnas.1509820112 (2015).

45 Kubota, Y. et al. A carbon nanotube tape for serial-section electron microscopy of brain ultrastructure. Nat Commun 9, 437, doi:10.1038/s41467-017-02768-7 (2018).

46 Lowe, D. G. Distinctive Image Features from Scale-Invariant Keypoints. International Journal of Computer Vision 60, 91–110, doi:10.1023/b:Visi.0000029664.99615.94 (2004).

47 Pekkurnaz, G., Trinidad, J. C., Wang, X., Kong, D. & Schwarz, T. L. Glucose regulates mitochondrial motility via Milton modification by O-GlcNAc transferase. Cell 158, 54–68, doi:10.1016/j.cell.2014.06.007 (2014).

48 Fiala, J. C. Reconstruct: a free editor for serial section microscopy. J Microsc 218, 52–61, doi:10.1111/j.1365-2818.2005.01466.x (2005).

49 Kerr, R. A. et al. Fast Monte Carlo Simulation Methods for Biological Reaction-Diffusion Systems in Solution and on Surfaces. SIAM J Sci Comput 30, 3126, doi:10.1137/070692017 (2008).

50 Stiles, J. R., Van Helden, D., Bartol, T. M., Jr., Salpeter, E. E. & Salpeter, M. M. Miniature endplate current rise times less than 100 microseconds from improved dual recordings can be modeled with passive acetylcholine diffusion from a synaptic vesicle. Proc Natl Acad Sci U S A 93, 5747–5752, doi:10.1073/pnas.93.12.5747 (1996).

51 Jorstad, A. et al. NeuroMorph: a toolset for the morphometric analysis and visualization of 3D models derived from electron microscopy image stacks. Neuroinformatics 13, 83–92, doi:10.1007/s12021-014-9242-5 (2015).

52 Asadulina, A., Conzelmann, M., Williams, E. A., Panzera, A. & Jekely, G. Object-based representation and analysis of light and electron microscopic volume data using Blender. BMC Bioinformatics 16, 229, doi:10.1186/s12859-015-0652-7 (2015).

